# Development and validation of a precise and accurate method to determine polyamine levels in cultured cells

**DOI:** 10.1101/2025.05.29.654479

**Authors:** Diana Cabrera, Mikel Pujana Vaquerizo, Belén Martínez La Osa, Amaia Zabala Letona, Arkaitz Carracedo, Juan Manuel Falcón, Sebastiaan van Liempd

## Abstract

We describe a robust, fast and accurate method for the quantification of intra-cellular concentrations of the polyamines, putrescine, spermidine and spermine in cultured cells. Hydrophilic interaction liquid chromatography in combination with Time-of-Flight mass spectrometry was used to obtain high resolution data for the analytes. Assay performance was determined with respect to chromatographic resolution, quantification, analyte recovery and matrix effects. Furthermore, assay variability was determined in a biological context. Based on these variability measurements, minimal detectable effects (MDEs) which would lead to significant differences in a null-hypothesis significance test, were calculated. As such, changes in spermine could be determined with the highest sensitivity with point estimates for the MDEs of 32% between-days and 10% within-days. For spermidine, these values were 38% between-days and 16% within-days. Finally, effects for putrescine were measured least sensitive with 43% between-days and 36% within-days. Finally, we employed the method to analyze the impact of polyamine synthesis pathway inhibition and cell culture conditions which are relevant aspects for the interpretation of the biological role of polyamines.

## Introduction

Polyamines (PAs) are biologically important metabolites. Due to their primary amino groups, they exist as cations under normal pH. They associate electrostatically with negatively charged macro-molecules like DNA and RNA. As such they play an important role in epigenetic processes. Furthermore, they are involved in redox balance and are associated with neuronal potassium channel activity. Overall, PAs are implicated in different diseases, such as cancer, immune response and neurological disorders [1].

There are three different biogenic PA species, namely putrescine (Put), spermidine (Spd) and spermine (Spm). Depending on the cell type, Put, Spd and Spm can have high intracellular concentrations (millimolar levels) and have well-stablished biological functions. Furthermore, polyamines can be acetylated to form various N-acetylated species. This process is considered a rate limiting step in polyamine metabolism [2]. We have focused this assay exclusively on Put, Spd and Spd.

Many methods are available for measuring biological PA concentrations. Most of these methods use some type of chromatography to separate the different PAs before detection with mass spectrometry [3]. Due to the highly polar nature of PAs, many such methods use derivatization [4] or ion-paring chromatography [5] to increase retention on conventional C18 column-chemistries. Derivatization also enables detection by UV or fluorescence. Furthermore, depending on the biological matrix, various types of homogenization and extractions have been described [3; 4; 6]. Some reported methods are very sensitive with a high dynamic range, generally depending on the type of mass spectrometer. Other assays focus on additional analytes, making them more versatile but less optimized for PAs [5]. More cumbersome are methods using derivatization reactions, which introduces more processing steps, and hence, more sources of variation. All these methods have their benefits and drawbacks and although many of these methods have good performance, we decided to establish our own to refine some of their limitations. Some methods described had long run times [7] or lacked proper separation. Moreover, some methods, like ion-paring chromatography or online SPE, were incompatible with our instrumentation. In any case, none of the available methods used the same equipment available in our lab or were not optimized for the desired biological matrix. Hence, the development of this assay to measure intracellular PA concentrations with ultra-performance liquid chromatography coupled Time-of-Flight mass spectrometry (UPLC-ToF-MS).

Chromatography was optimized with respect to carry-over and chromatographic resolution. Furthermore, we have determined dynamic range and lowest limit of quantification (LLOQ) in solution. All other development steps were carried out in a biological context, meaning that we used biological matrix in the various optimization experiments. Extraction and injection liquids were optimized with respect to carry-over, signal intensity and recovery. Furthermore, we describe a method for the determination of ion-suppression due to matrix effects using spiked quality control samples. By using cell extracts to determine inter-, intra-day and measurement variability, it was possible to calculate minimal detectable effects (MDEs) in a realistic, biological scenario using Bayesian estimates.

We established a simple, fast, sensitive and well-defined analytical method to measure intracellular concentrations of Put, Spd and Spm in cultured cells. A simple biphasic extraction procedure was used to clean up samples and to extract and concentrate analytes. High-resolution hydrophilic interaction liquid chromatography (HILIC) coupled to ToF-MS was used for chemical analysis.

We used our newly established method to determine the effect of inhibition of the enzyme S-adenosylmethionine decarboxylase (SAMDC) by SAM486A on PA pools using ^13^C_5_-methionine as a precursor to track ^13^C labelled carbons. Since SAM486A is a well-known inhibitor and its effects are well documented, this experiment served as a positive control of the performance of the analytical method. Finally, we used the method to measure intracellular analyte concentrations for DU145 cells that were incubated with different culture media conditions.

## Methods

### Materials

Optima® LC/MS grade water (water), methanol (MeOH), acetonitrile (ACN) and formic acid (FA) were purchased from Fischer Scientific (Loughborough, UK). Ammonium formate (AmFo) and acetic acid (AA) were purchased from Honeywell/Fluka (Seelze, Germany). Chloroform as well as putrescine dihydrochloride, spermidine trihydrochloride and spermine tetrahydrochloride were purchased from Sigma-Aldrich (Steinheim, Germany). L-Methionine-^13^C_5_ was purchased from Cambridge Isotope Laboratories. SAM486A was kindly provided by Novartis (referenced in [8]). Safe-Lock™ 1.5 mL microtubes were purchased from Eppendorf (Hamburg, Germany). Polypropylene Acquity UPLC 700 µL round 96-well-plates were purchased from Waters (Manchester, UK) as well as silicon/PTFE treated pre-slit seals.

### Solutions

Optimized extraction liquid (EL) consisted of 10 mM AA in 50% (v/v) MeOH/water. Optimized resuspension liquid (RL) consisted of 39.8% water, 60% ACN and 0.2% FA (v/v/v).

### Reagents for cell cultures

Human prostate carcinoma cell line DU145 (ACC261) was used and purchased from Leibniz-Institut DSMZ (Deutsche Sammlung von Mikroorganismen und Zellkulturen GmbH), which provided authentication certificate, and were tested for mycoplasma every two weeks. Cells were cultured in Dulbecco’s modified eagle medium (DMEM, Gibco^Tm^41966-029 and Penicillin/ Streptomycin (Gibco^Tm^15140-122) (0.1%) with 10% of Fetal Bovine Serum (FBS) (Gibco^Tm^ 10270-106) or Dialyzed FBS (Thermo Fisher Scientific A3382001). Human Plasma Like-Medium (Thermo Scientific A4899101) was also used as a third approach.

### Cellular assay

For the assay variability test, DU145 cells were seeded in four 6-well plates. For each measurement day a single plate was used. To test culture medium conditions, DU145 cells were seeded in 6-well plates in four independent experiments and with 3 technical replicates per experiment. After 6 hours, the cells were attached, and we changed the media to culture them in the three different media (DMEM complete; DMEM dialyzed serum; HPLM) for 48h when they reached a confluence of 80% in the well. Finally, cells were washed three times with abundant phosphate buffered saline (PBS, Gibco^Tm^14190-034) and snap frozen in liquid nitrogen before they were stored at −80 °C. For inhibition experiments SAM486A was added to DMEM complete medium in a concentration of 500 nM. Moreover, to label Spm and Spd, uniformly ^13^C-labelled methionine (^13^C_5_-methionine) was added to DMEM complete medium to obtain a final concentration of 0.2 mM labelled methionine and a total methionine concentration (labelled and unlabelled) of 0.4 mM. For inhibition experiments after 42h, medium was changed to the medium containing SAM486A and ^13^C_5_-methionine and cells were further incubated for 6 hours.

### Optimalization of cell extractions

To optimize the extraction procedure, two monophasic and one biphasic extraction method were tested. For the monophasic extractions the extraction liquid contained either 50% (M50) or 80% MeOH (M80) in water (v/v). Both solvent mixtures contained 10 mM AA. The extraction liquid of the biphasic method (BP) contained 10 mM AA in 50% MeOH in water (v/v) for the aqueous phase and chloroform for the organic phase.

Cells were extracted by putting the six-well culture plates on ice, adding 500 µL ice-cold extraction liquid to the wells and letting these rest for 15 minutes. Subsequently, the cells were scraped of the bottom of the wells and 400 µL of this suspension was transferred to 1.5 mL microtubes. For the biphasic extraction 400 µL chloroform was added to this homogenate.

The extractions were agitated for 30 minutes at 1400 rpm and 4 °C and the microtubes were centrifuged for 30 minutes at 14000 rpm (∼13200 g) and 4 °C. Next, 250 µL of the supernatant was transferred to a clean microtube and evaporated to dryness. For the biphasic extraction 250 µL of the aqueous layer was used (top layer). Pellets were resuspended in 150 µL 60% ACN/40% water (v/v) and measured.

### Analyte recovery

Recovery was determined by using samples that were either spiked before (pre-spike) or after extraction (post-spike) and non-spike samples. Extractions were carried out with the biphasic extraction method. For pre-spiked samples, either 100 nM or 1 µM of the three analytes were spiked in EL which was then used to extract cells. For post-spike samples, analytes were spiked in RL at the same concentrations. Non-spiked samples were worked up without spiking analytes in either liquid. Volumes taken from the aqueous phase for evaporation and for resuspension of the resulting pellets were both 200 µL.

### Sample preparation

For variability tests and method application, the biphasic extraction method was used with a slight modification in the transferred volumes. After centrifugation, from each extraction 280 µL of the aqueous phase was transferred to a clean 1.5 mL microtube and from this tube 30 µL was transferred to a second 1.5 mL microtube. Thus, each cell extraction results in two microtubes containing 250 µL which content was used for the analytic sample and 30 µL aqueous phase which was used for quality controls (QCs).

The aqueous phases were evaporated in a speed vac to dryness in two batches. The 250 µL samples took about 3 hours to fully evaporate and 30 µL samples took about 30 minutes. The pellet of each analytic sample was resuspended in 150 µL RL. The pellets of the QC samples were resuspended sequentially in a single volume which depended on the number of extractions. For each extraction, 9 µL RL was added, *e.g.* when 30 extractions were analyzed, a single volume of 30 × 9 µL = 270 µL was used to resuspend all QC samples. This was done by adding the single volume to an evaporated QC sample, resuspending the pellet and transferring it to the next QC sample *etc*. until all QC samples were resuspended in the single volume.

The QC sample was then split into two equal parts. To one part, an equal volume of EL was added. These were used for initialization QCs (QC_init_) and normal QC samples (QC_0_). To the second part, the same volume of 10 µM analyte-spiked EL was added (QC_spike_). Following the above example, with 30 extractions, two aliquots of 135 µL were created. To one of these, 135 µL EL was added and to the other 135 µL 10 µM spiked EL was added. This to ensure that the matrix in the QCs was the same as in the average analytical sample. The QC_spike_, which had added analyte concentrations of 5 µM, could be used to estimate the matrix effects affecting signal strength.

After resuspension, all samples were centrifuged again for 15 minutes at 14000 rpm (∼13200 g) and 4 °C. Finally, 130 µL of the analytical sample and the total volumes minus 20 µL of the QC samples were transferred to a 96-wells plate for injection on the LCMS system.

### Analytical standards and calibration samples

Separate analytical standards of Put, Spd and Spm with a concentration of 100 mM were made in water and stored in 4 mL glass tubes at −20 °C for further use. These standards were used to make a pooled solution of 1 mM in water which was stored in a microtube at −20 °C for further use. This pooled solution was then used for making calibration curves and QC_spike_ solutions.

To make the calibration curve, 100 µL pooled 1 mM analyte solution was added to 300 µL water containing 0.66% (v/v) FA and subsequently 600 µL ACN was added. This resulted in a 100 µM solution in RL. This 100 µM solution was diluted ten times in RL to obtain 2 mL 10 µM analyte solution. This solution was used to spike the QC_spike_ sample and for the highest point of the calibration curve.

The dilution series for the curve was as follows: 10 µM was diluted two times in RL to 5 µM. The 5 µM solution was diluted another two times to 2.5 µM. The 10 µM, 5µM and 2.5µM solutions were ten times diluted to obtain 1 µM, 0.5 µM and 0.25 µM. The resulting solutions were again ten times diluted to 0.1 µM, 0.05 µM and 0.025 µM. The 0.025 µM solution was diluted two times, four times to get the following concentration range: 12.5 nM, 6.3 nM, 3.1 nM and 1.6 nM. Thus, the curve contained 14 points including a RL blank.

### Sample sequence

The injection sequence used for the variability tests and method application started with the 14-point calibration curve, followed by four QC_init_ samples to equilibrate the column. Next a QC_0_ and a QC_spike_ were injected followed by the analytical samples. QC_0_ and QC_spike_ were injected at regular intervals between the samples and at the end of the sample sequence. The QC interval depended on the number samples and was around 6 to 8 samples. After the final QC set, another calibration curve was injected. The analytical samples were injected in duplicate or in triplicate and in a randomized order.

### Liquid chromatography

An Acquity I-Class Plus UPLC system (Waters) was used for chromatographic separations. The system was equipped with a binary solvent manager, a two-position sample manager, a ten-position sample organizer and a column heater. A 12.5” HPS column active preheater was used to connect the injection valve with the column inlet.

LC optimalization was carried out by testing different mobile phases and gradients as described in Table 1) and Table 2). The optimized mobile phases consisted of aqueous phase (A) containing water, 0.25% FA (v/v) and 2.5 mM AmFo and an organic phase (B) containing 9.75% water, 0.25% FA, 90% ACN (v/v/v) and 2.5 mM AmFo. To prepare the B phase, FA and AmFo were first mixed with water and then ACN was added.

**Table 1.**
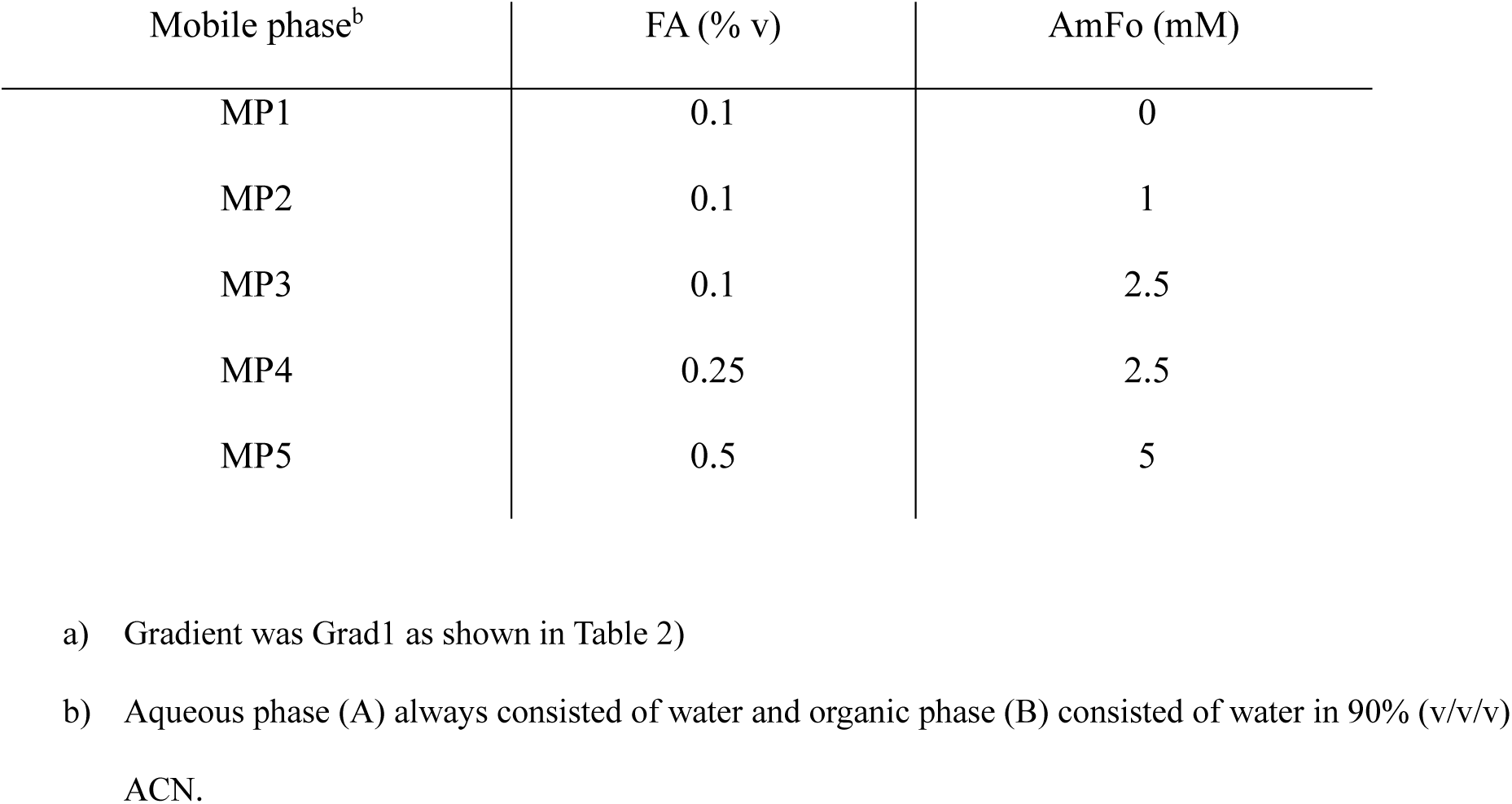
Mobile phase optimalization. Formic acid (FA) and ammonium formate (AmFo) content in both aqueous and organic mobile phases. Same elution gradient^a^ was used for all tests.

**Table 2.** LC optimalization. Gradients tested.

| Gradient | %A<br>start<br>gradient | Time<br>gradient<br>(min) | %A end<br>gradient | Time<br>strong<br>(min) | %A<br>end<br>strong | Time<br>gradient<br>(min) | %A<br>Start<br>equilibration | Equilibration<br>time<br>(min) |
| --- | --- | --- | --- | --- | --- | --- | --- | --- |
| Grad1 | 50 | 2 | 100 | 2.4 | 100 | 0.1 | 50 | 1 |
| Grad2 | 40 | 2 | 100 | 2.4 | 100 | 0.1 | 40 | 1 |
| Grad3 | 50 | 3 | 100 | 1.4 | 100 | 0.1 | 50 | 1 |
| Grad4 | 40 | 3 | 100 | 1.4 | 100 | 0.1 | 40 | 1 |
| Grad5 | 45 | 3 | 100 | 1.4 | 100 | 0.1 | 45 | 1 |
| Grad6 | 50 | 3.5 | 100 | 0.9 | 100 | 0.1 | 50 | 1 |
| Grad7 | 52 | 3.5 | 100 | 0.9 | 100 | 0.1 | 52 | 1 |

Separation was carried out on an Atlantis™ Premier BEH Z-HILIC 1.7 µm 2.1×100 mm column. The column was thermostated at 40 °C. The linear gradient was as follows: 52%A to 100%A in 3.5 minutes, constant at 100%A for 0.9 minutes, back to 52%A in 0.1 minutes and constant at 52%A for 1 minute for column equilibration. The flow rate was set to 400 µL/min.

### Mass spectrometry

Mass spectrometric analysis was performed on a SYNAPT G2S instrument (Waters). The instrument was used in full-scan (50-1200 Da) positive electrospray ionization mode. The optimized source settings were as follows: the capillary voltage was set to 200 V, sampling cone voltage to 25 V, source offset voltage to 80 V. Source temperature was 120 °C and desolvation temperature was 450 °C. The cone gas flow was 5 L/h, nebulizer gas flow was 6 bar and desolvation gas flow was 600 L/h. All gas flows were nitrogen. Scan rate was set to 0.2 seconds. Accurate mass was assured by infusing a lock-mass (Leu-Enk, 100 ppb in 49.9% water, 50% ACN, 0.1%FA (v/v/v), *m/z* = 556.2771) at 10 µL/minute every 60 seconds for 0.5 seconds.

### Data analysis

Extracted ion chromatograms (EICs) for the protonated forms of the analytes, Put (*m/z* = 89.1079), Spd (*m/z* = 146.1657), ^13^C_3_-Spd (*m/z* = 149.1758), Spm (*m/z* = 203.2236), ^13^C_3_-Spm (*m/z* = 206.2336) and ^13^C_6_-Spm (*m/z* = 209.2437) were extracted in a 20 mDa window and integrated with TargetLynx software (Waters). EICs were smoothed by taking a two-point average for two iterations. Peak areas were used for further analysis. To determine the fraction of ^13^C-labelled PAs, the fraction of natural occurring ^13^C was subtracted from the signal in the mass channels of the ^13^C_3_ (Spd/Spm) and ^13^C_6_ (Spm) species. All calculations in this manuscript were performed in R (version 4.4.1).

Signal data was pretreated to adjust for biases during sample work-up and analysis. Analysis scripts are available (Supplemental information). First, analyte signals were adjusted with median fold-change (MFC) normalization [9; 10]. In short, 1) LCMS features (*i.e.* retention time–*m/z* pairs) were collected for each sample, including QCs. 2) A reference sample was determined based on the highest number of features. 3) For each sample, each feature was scaled on the value of the reference sample, obtaining fold-change value for each feature over all included samples. 4) The median of the fold-change values for each sample was calculated to obtain the MFC normalization factor for each sample. 5) Analyte signals were divided by the MFC factor. This method adjusts for factors like differences in biological material or variation in extraction between samples and altering detector response during the run.

The remaining variation in analyte signals was determined and possibly corrected with QC_0_ values. 1) The QC_0_ signals per analyte were normalized on the first QC_0_. 2) For each analyte, a polynomial of the first degree was fitted through the normalized signals by using the injection number as the independent variable. 3) The sample signals were corrected by dividing the signal by the value returned by the polynomial, based on the injection numbers of the samples. If either MFC-normalization or QC-correction did not lead to a median drop in variation between replicate injections, the correction was not used and instead the raw or single-corrected signals were used for further calculations.

Qualification was done by using the following algorithm. 1) The median signal in the sample space was determined. 2) The curve concentration with the closest signal to this median signal was chosen as focus concentration. 3) All 4-point log-transformed linear models containing this focus concentration were made, using both curves. 4) Deviation from the nominal curve concentrations were determined for each model. 5) The model for which the least calculated concentrations deviated more than 15% from their nominal values was chosen for sample quantification.

To show signal behavior over the entire calibration curve, curves were fitted with log-transformed linear regression to account for the non-linear response of the detector. First, calibration points were excluded that were lower than 12 times the baseline noise (lowest limit of quantification, LLOQ). For the remaining points the concentration (*x*) and signal (*y*) values were log transformed. Next, a linear model was fitted to the transformed data. This model can be interpreted as follows: the intercept (*α*) and slope (*β*) of the linear model can be used on the non-transformed data as a power-law relationship *y = e^α^ x^β^*. To determine a concentration from a signal, the following relation can be used: *x = (y/e^α^)^1/β^*. Both curves were used to create the linear model.

After obtaining accurate (adjusted) signal and concentration data, various assay metrics were determined. These metrics included extraction efficiency, recovery, signal attenuation due to matrix effects and assay variability from biological and instrumental sources. The final method was used to determine cell concentrations of the analyte for different culture media types. For statistical analysis we used Bayesian multilevel models. Bayesian models combine prior information and the likelihood of the data to generate a posterior distribution of the model parameters. The parameter estimates can then be used to generate posterior distributions of the desired metrics. These models provide the flexibility needed to handle complex data and return full probability distributions for parameters, offering a natural way to quantify uncertainty.

All models were given a normal likelihood, truncated at 0, to assure that the posterior samples were strictly positive as is the case for signal or concentration data. Priors for fixed and random effects were normal distributions and priors for standard deviations were exponential distributions. Random effects were assigned where possible for repeated measures, like sample replicates. Experimental groups were treated as fixed effects. Furthermore, the model used to calculate recoveries assured that estimated means for the no-spiked group should always be smaller than that of the pre-spiked group and that of the pre-spiked group should be smaller than that of the post-spiked group, *i.e.* µ_0_ < µ_pre_ < µ_post_. Bayesian models were written in the probabilistic programming language Stan (version 2.32.2). Posterior parameter estimates for treatment groups, obtained from the various models, were used for further calculations. The code for the various models is available in the Supplementary Information.

Extraction efficiency was estimated simply by comparing the group estimates for the different extraction groups. The same was done to estimate cell concentrations for different culture media types. Recovery was determined by first estimating the expected signal (*E[Signal]*) for the pre-spiked, post-spiked, and non-spiked sample groups. Next, baseline-adjusted estimates were obtained by subtracting the expected signal of the non-spiked group from both the pre-spiked and post-spiked estimates. Recovery was then calculated as the ratio of the baseline-adjusted pre-spiked estimate to the baseline-adjusted post-spiked estimate. Mathematically, this is defined as: *R=(E[Signal_pre_] - E[Signal_0_]) / (E[Signal_post_] - E[Signal_0_])*, where *E[Signal_pre_]*, *E[Signal_post_]*, and *E[Signal_0_]* represent the expected signal for the pre-spiked, post-spiked, and non-spiked groups, respectively. Attenuation due to matrix effects was determined similarly. By dividing the posterior means of the QC_0_-subtracted QC_spike_ group (spike level at 5 µM) by the posterior mean of the 5 µM signal from the calibration curves in solution, attenuation was determined. The calculation was as follows: *A = (Signal_QCspike_ – Signal_QC0_)/Signal_curve_*. To determine inter-day, intra-day and measurement variation, the estimates of the relative standard deviations (equivalent to coefficients of variations, %CV) for run, sample and likelihood were used. The relative standard deviations were used to calculate minimal detectable effect (MDE) per analyte. The standard error was calculated using the square root of the sum of variances, assuming n=6. The MDE was then determined by multiplying the standard error with the critical t-value for α=0 (two-sided, n=6).

## Results and Discussion

The method was optimized for chromatography and extraction efficiency until acceptable performance was achieved taking in account chromatographic resolution, signal intensity and carry-over. We tested five mobile phases, seven LC gradients and three extraction methods. Importantly the assay was validated in a biological context. This means that recovery, matrix effects and variability were all determined in the target matrix rather than in BSA solutions or solvent as is the case in some assay validations, for example [6; 7]. This gives a more realistic idea of the assay performance. Finally, we applied the method to determine differences in PA levels for cell cultures, using different culture media conditions.

### LC method optimalization

Polyamines (PAs) are extremely hydrophilic with logP-values for Put, Spd and Spm of respectively −0.466, −0.504 and −0.543. Therefore, we tested two hydrophilic column chemistries namely BEH AMIDE and BEH Z-HILIC before settling on the latter one. Although the AMIDE column did retain and separated the PAs sufficiently well, the gradients and mobile phases were more complex, and the peak shapes were worse compared to the Z-HILIC column (results not shown).

For optimalization of the Z-HILIC method we started with optimizing the mobile phase. For this we increased FA and AmFo content in both aqueous and organic phases. This resulted in increased resolution and intensity (Fig. 1). Specifically, higher AmFo content resulted in increased resolution and sharper peaks (*i.e.* MP2 *vs.* MP3) while increased FA content resulted in increased signal strength (*i.e.* MP3 *vs.* MP4) (Fig. 1). The highest chromatographic resolutions (Table 3) were obtained with MP5 which contained the highest amount of AmFo.

**Figure 1.**
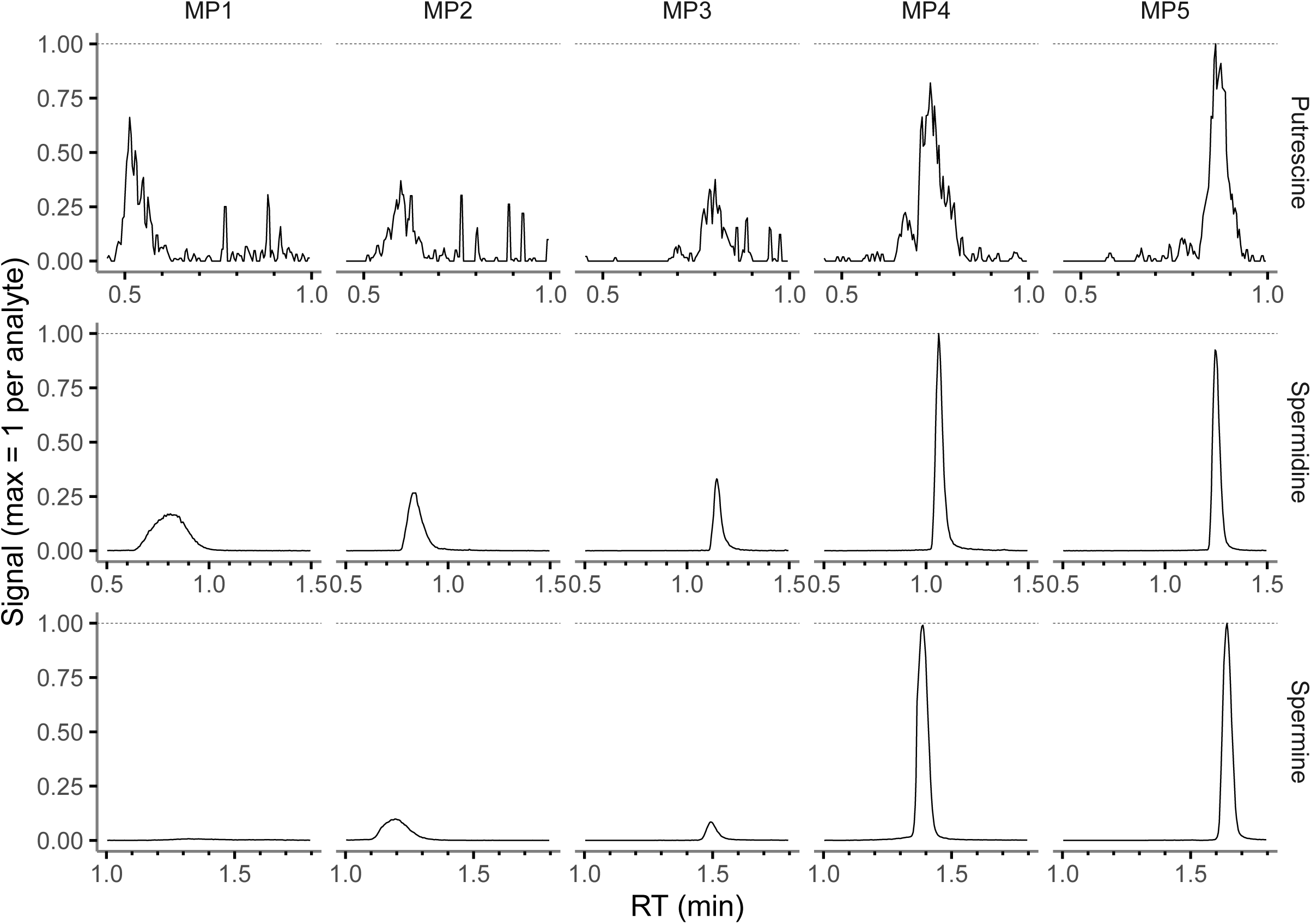
Mobile phase optimalization using 10 µM analyte mix and Grad1. The signals were smoothed by averaging over each two adjacent values. Subsequently smoothed values were scaled on the maximum value per analyte. Mobile phases and gradient are described in Table 1 and 2.

**Table 3.** Chromatographic resolutions^a^ for tested mobile phases and gradients.

| Optimalization | Gradient | Mobile phase | R1 | R2 |
| --- | --- | --- | --- | --- |
| Mobile phase | Grad1 | MP1 | 2.7 | 2.3 |
| Mobile phase | Grad1 | MP2 | 3.7 | 4.4 |
| Mobile phase | Grad1 | MP3 | 7.9 | 9.3 |
| Mobile phase | Grad1 | MP4 | 6.6 | 9.3 |
| Mobile phase | Grad1 | MP5 | 9.7 | 12.1 |
| Gradient | Grad1 | MP4 | 7.2 | 8.8 |
| Gradient | Grad2 | MP4 | 11.9 | 10.0 |
| Gradient | Grad3 | MP4 | 7.5 | 10.9 |
| Gradient | Grad4 | MP4 | 13.8 | 11.9 |
| Gradient | Grad5 | MP4 | 10.4 | 12.3 |
| Gradient | Grad6 | MP4 | 8.7 | 12.1 |
| Gradient | Grad7 | MP4 | 11.7 | 12.5 |

a) R1: resolution between putrescine and spermidine; R2: resolution between spermidine and spermine.

The increase of FA from 0.1% (MP3) to 0.25% (MP4) resulted in approximately 2-times (Put), 3-times (Spd) and 10-times (Spm) increase of signal strength. Increase to 0.5% FA (MP5) resulted in only slightly increased (Put) or decreased (Spm) signal strength.

Although MP5 gave the best resolution we decided to use MP4 for further optimalization. This choice was made to avoid the risk of crystallization of AmFo due to its high concentration (5 mM) in the organic phase. Moreover, higher AmFo concentrations could lead to faster contamination of the ion source and lower column-life. On the other hand, the difference in signal strength and resolution were not critically worse than with MP4.

After choosing a mobile phase, the gradient was optimized. For this we varied the initial mobile phase proportions and the gradient time. All gradients showed excellent resolutions (Table 3) with Grad4 and Grad7 as optimal. However, we found that gradients starting below 50% A had some carry-over (not shown) while gradients starting at 50% A or higher showed clean blank injections after injection of a high (10 µM) analyte concentration. Moreover, increasing gradient time from 2 to 3 minutes increased resolution even further. Therefore, for subsequent analysis we used Grad7, which starts at 52% A and has a gradient time of 3 minutes.

Another source of carry-over was the absence of FA in the injection liquid. Signals in blank injections after injection of a 10 µM PA mix sample without FA showed signals for Spd and Spm, equivalent to nM levels (Fig. 3). These signals disappeared with FA concentrations of 0.1% or higher in the injection liquid. Therefore, the injection liquid should at least contain 0.1% FA. However, when preparing and measuring samples over various days, we found that 0.1% FA injection liquid was not stable during prolonged storage (> 3 days), since carry-over increased over time. This is probably due to the evaporation of FA. Therefore, we increased FA content to 0.2% which resulted in an injection liquid that was stable for at least a week. Injection liquid containing 39.8% water, 60% ACN and 0.2% FA (v/v/v) was used for further experiments.

**Figure 2.**
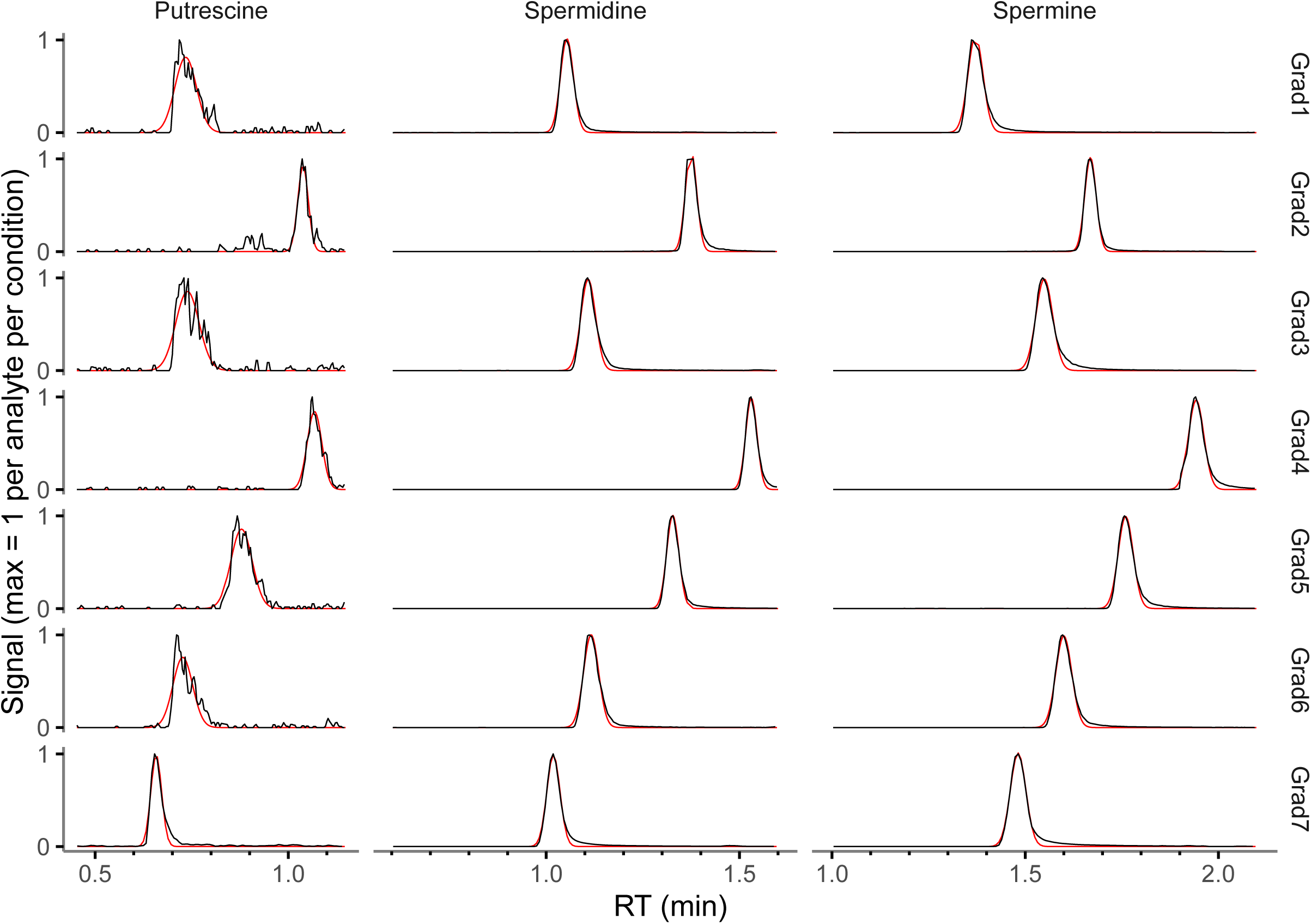
Gradient optimalizations using 10 µM analyte mix and MP4. The signals (black lines) were smoothed by averaging over each two adjacent values. Subsequently smoothed values were scaled on the maximum value per analyte and condition. A gaussian was fitted on the data (red lines). Mobile phase and gradients are described in Table 1 and 2.

**Figure 3.**
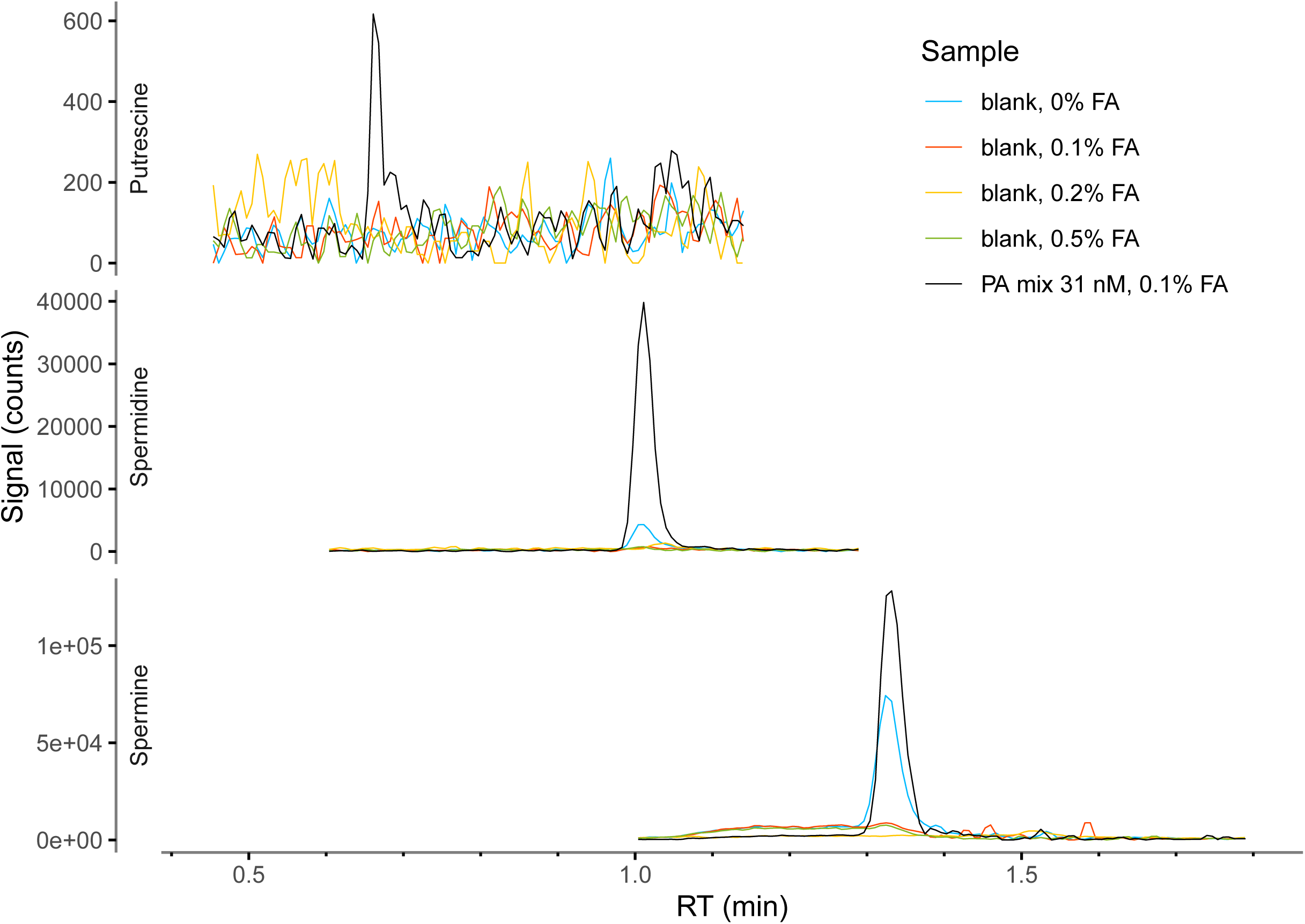
Carry over in blank samples, which were directly injected after 10 µM mix, given different concentrations of formic acid in the injection liquid (blue: 0%FA, orange: 0.1% FA, yellow: 0.5% FA). Also shown (black) is a 31 nM mix sample for comparison. The signals were smoothed by averaging over each two adjacent values.

### Extraction efficiency

To optimize the extraction of the targeted analytes from cells, we tested three extraction methods. Given that the volumes subjected to evaporation and subsequent resuspension were the same for each method, the method resulting in the highest signals would be optimal. We found that the M80 method performed worse for all three analytes (Fig. 4A). This might be due to the highly polar nature of the analytes which could cause the analytes to stick to matrix components rather than to dissolve in a high concentration of methanol. The M50 method showed similar performance for Put and Spd but was significantly worse for Spm. This might be due to the same reason as given for M80. However, since Put and Spd are less polar than Spm, the partition equilibrium of Put and Spd over matrix components and solvent may lay more towards solvent in M50 than that of Spd. It could be argued that lowering the MeOH content even further would result in a higher extraction efficiency. However, from experience we found that MeOH content lower than 50% resulted in worse protein precipitation which led to problems such as clogging of the chromatographic column.

**Figure 4.**
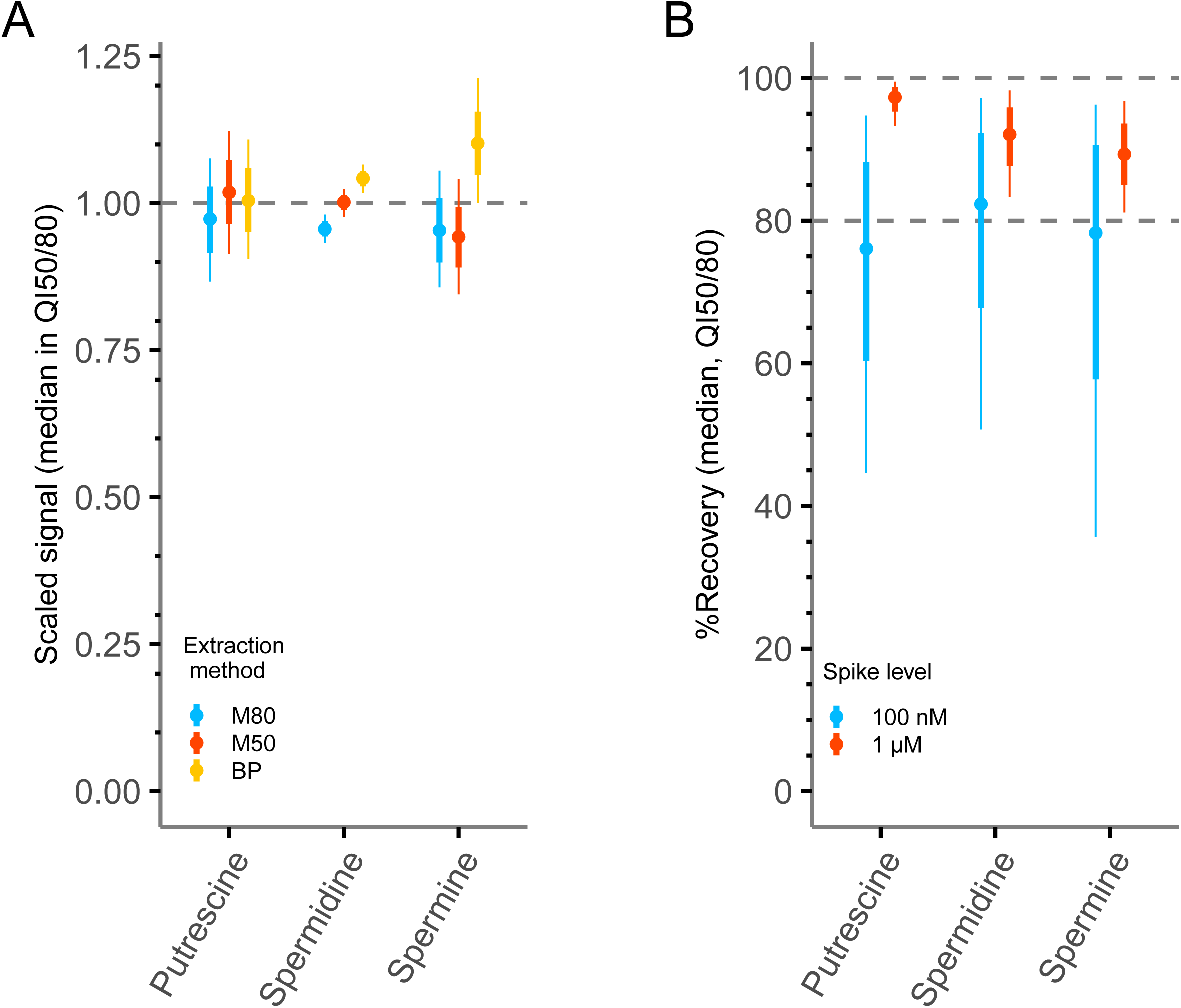
Extraction efficiency for tested extraction methods (A) and recoveries (B). For visualizations of extraction efficiencies, signals were scaled on the overall data-mean per metabolite (*e.g.* mean = 1). Values higher than 1 indicate higher signals than the mean for that extraction and *vice versa*, indicating extraction efficiency. For example, for spermine BP (∼1.3) performs better than M80 or M50 (∼0.8). Analyte recovery from cell homogenates for two spike levels. Recoveries for 1 µM spikes were 90% or higher. Recoveries for spikes at 100 nM could not be determined accurately due to the high background signals in the homogenates. For both extraction and recovery plots, posterior median values within 50% and 80% quantile intervals from Bayesian models (Supp1) were used. M80: 80% MeOH/20% water, M50: 50% MeOH/water, BP: biphasic: (50% MeOH/water):(chloroform).

The best performing extraction for all analytes was the biphasic one (BP). This could be because part of the MeOH dissolves in the chloroform which leaves the aqueous phase more polar and thus dissolves the polyamines better. Moreover, many apolar components were transferred to the organic phase, leaving the sample cleaner.

### Analyte recovery

After optimizing the extraction liquid, the amount of recovered analyte was determined for two spike levels, *i.e.* 100 nM and 1 µM. Therefore, cell homogenates were spiked before and after extraction at two levels. Non-spiked samples were then used to subtract background analyte signals. We found good recovery for 1 µM spike levels (Fig. 4B, Table 4). Recovery for Put was 97% (95%-99%), for Spd 92% (88%-96%) and for Spm 89% (85%-94%). The recovery for 100 nM could not be determined accurately because of the high endogenous signal, relative to the spiked-in concentrations. Although model estimates suggest that recoveries for 100 nM spikes are somewhat lower. However, since the endogenous concentrations were close to the 1 µM spikes, the recovery at 1 µM should be indicative for regular cell extractions.

**Table 4.** Assay metrics including recovery, signal attenuation. Values are given as the posterior median with the 50% quantile interval between parentheses.

| Analyte | Recovery (%)<sup>a</sup> |  | Attenuation<sup>b</sup> |
| --- | --- | --- | --- |
| | 100 nM | 1 μM | |
| Putrescine | 76 (60-88) | 97 (95-99) | 0.35 (0.32-0.38) |
| Spermidine | 82 (68-92) | 92 (88-96) | 0.59 (0.55-0.68) |
| Spermine | 78 (58-91) | 89 (85-94) | 0.87 (0.83-0.91) |

a) Recovery for two spike levels, *i.e.* 100 nM and 100 μM.
b) Attenuation was defined as the factor by which the signal in the biological matrix was attenuated with respect to the signal in solution.

### Quantification

To illustrate signal behavior as function of nominal curve concentrations, calibration curves were analyzed over the total concentration range. Note that final quantification was done with a legacy method that only uses the part of the curves that are most representative for sample signals. It was found that by using our legacy quantification method, the inter-day assay variability was significantly lower (results not shown).

Because older ToF instruments show strong non-linear behavior over extended concentration ranges, curve fitting is not straight forward. Even when a double log-transform was done, there still was some non-linear behavior left. The parts before and after 0.5 µM have different slopes (Fig. 5A). Therefore, two curves could be fitted for the lower and upper part with a common point at 0.5 µM. Considering adjusted R^2^ values, which did all exceed 0.98, all fits were good. Mean deviations from nominal values for all three analytes, as measured over 5 runs, were all within ±15%. This indicates that the assay has high accuracy [11]. The standard error of the mean-intervals (SEM) for Spd and Spm also mostly did not surpass this threshold, the Put SEM -intervals are in most cases somewhat higher indicating that the assay is less accurate for this metabolite. This is not surprising given that the signal-to-noise ratios for Put were relatively low, compared to Spd and Spm.

**Figure 5.**
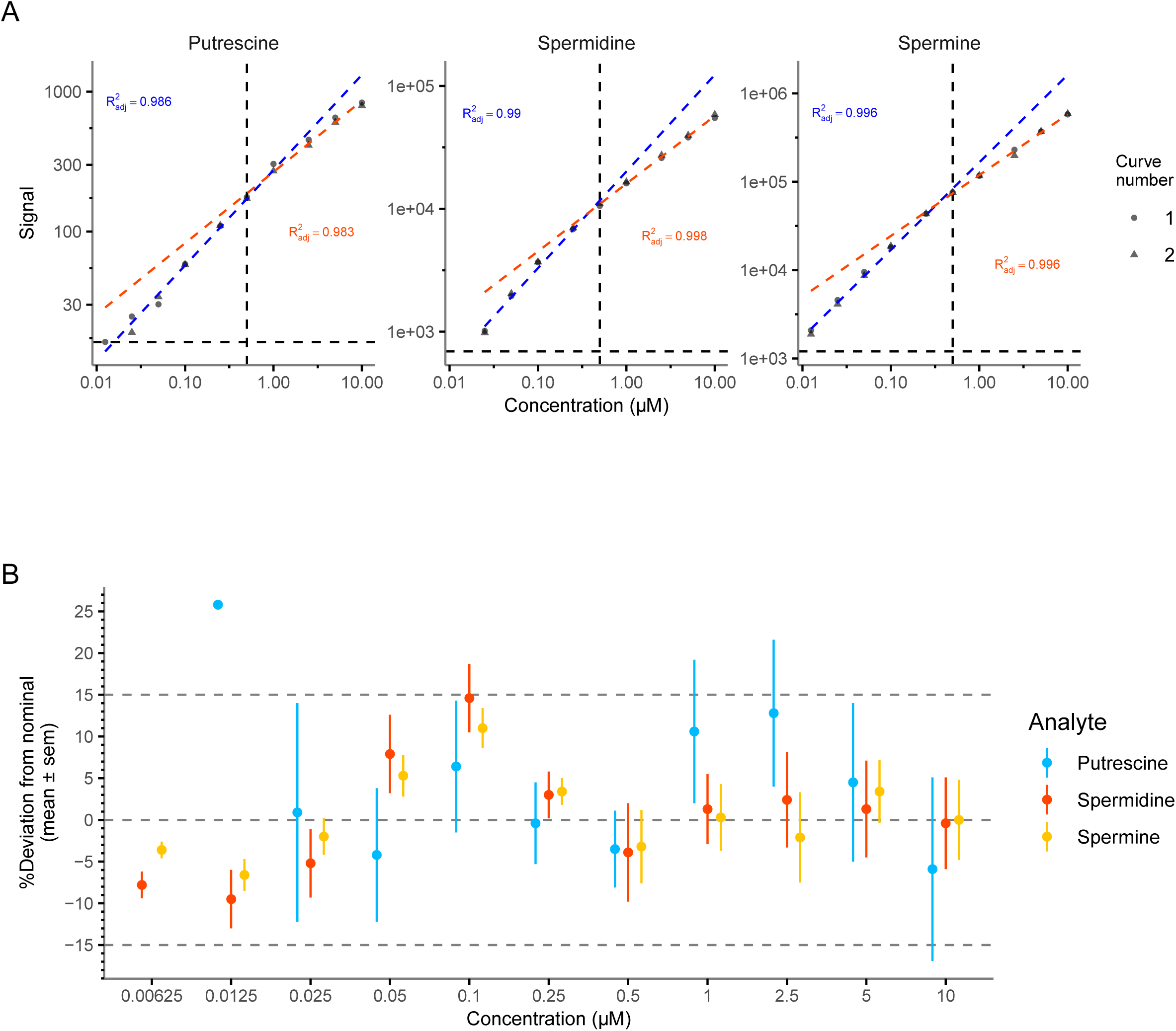
Calibration curves for the measured analytes for a single run (A) and deviations from nominal concentrations for all runs (B). Curves were plotted on a log-log scale for better visualization of signals over the total concentration range. Curves were fitted with log-transformed linear regression. For each analyte two curves were fitted for the lower (blue) and the upper part (orange). The common point for both parts was 0.5 µM (vertical dotted line). The horizontal dotted line represents the LLOQ. Adjusted R^2^ values for both lower and upper curves are shown. Deviations from nominal concentrations were are based on 5 calibration runs. Deviations are given as percentages within the standard of the mean (SEM).

For 2 µL injections, the LLOQs for Put and Spd were around 10 nM while that of Spm was around 3 nM. Since curve samples were measured in solution no background signals had to be extracted which increased the sensitivity of the assay. However, a disadvantage of measuring calibration samples in the absence of matrix is that ion-suppression in the biological samples is not accounted for. However, by using spiked and non-spiked QC samples an estimation of matrix effects could be made as explained in the next section. The sample concentrations could then be adjusted accordingly.

In any case, the LLOQ levels were more than sufficient to determine concentrations for the number of cells that were used for variability tests, *i.e.* 500k cells per sample. For these samples, vial concentrations for the analytes, *i.e.* 650 nM for Put, 1.8 µM for Spd and 250 nM for Spm, lay well above their LLOQs.

### Signal attenuation

Matrix components can cause analyte signals to change in intensity with respect to signals measured in pure solvent solutions. Normally, matrix effects lead to a decrease in ionization efficiency and hence a decrease in signal strength. This could be a problem when a calibration curve is prepared in solution as in this study. In the case of signal attenuation in matrix, this could lead to lower apparent concentrations.

There are various ways to correct for these effects. Obviously, a curve could be prepared in matrix. However, as explained, this would lead to an increase in the quantification limit due to the high endogenous background. Another way to correct is to add an internal standard to the resuspension liquid. This internal standard could be a non-endogenous compound with similar chemical properties or a stable isotope-labelled analyte. Both these methods have drawbacks. We added cadaverine (Cad), a chemically similar, non-endogenous compound, as an internal standard. However, this surprisingly led to severe suppression of the Put signal. This was presumably associated with the fact that Cad and Put coeluted and that the concentration of the spiked Cad (10 µM) was high. Moreover, as we will see later, adding a biosimilar compound will not be sufficient to correct for all analyte attenuations. As for stable-labelled analytes, at the time of development, besides being very expensive these compounds were only available in small quantities, which made it very challenging to make accurate stock solutions. Additionally, if metabolic flux experiments are carried out using stable isotope tracers, these internal standards could interfere with labelled analyte signals.

Therefore, we chose to correct for signal attenuation by injecting a spiked QC sample after each normal QC. Then, the difference in signal between these two samples was compared to a sample of the same analyte concentration but in solution. We found that attenuations varied strongly between analytes. The signal of Put in matrix was 0.35-times the signal in solution, for Spd it was 0.59-times and for Spm 0.87-times (Table 4). This also shows that a single internal standard is not enough to correct for the various attenuation effects.

However, the measured concentrations in this study were not corrected for these effects because unfortunately we did not include this method in some of the validation runs. Therefore, if this method was applied to only some of the runs, a comparison over all runs (necessary to estimate assay variability) would not be possible. Notwithstanding, the authors do recommend that this correction should be made when the method is applied in future studies.

### Assay variability

We determined sources of variation by measuring analyte concentrations in cell cultures over four days (Fig. 6). This validation process differs somewhat from a conventional biological assay validation as described by the FDA guidelines. The guidelines recommend using the variability in spiked QC samples to estimate assay stability. This gives a good estimation of technical variability. The method to estimate assay stability also takes into account biological variation and thus is more conservative. Moreover, we were able to calculate minimal detectable effects as expected in future measurements using biological samples.

**Figure 6.**
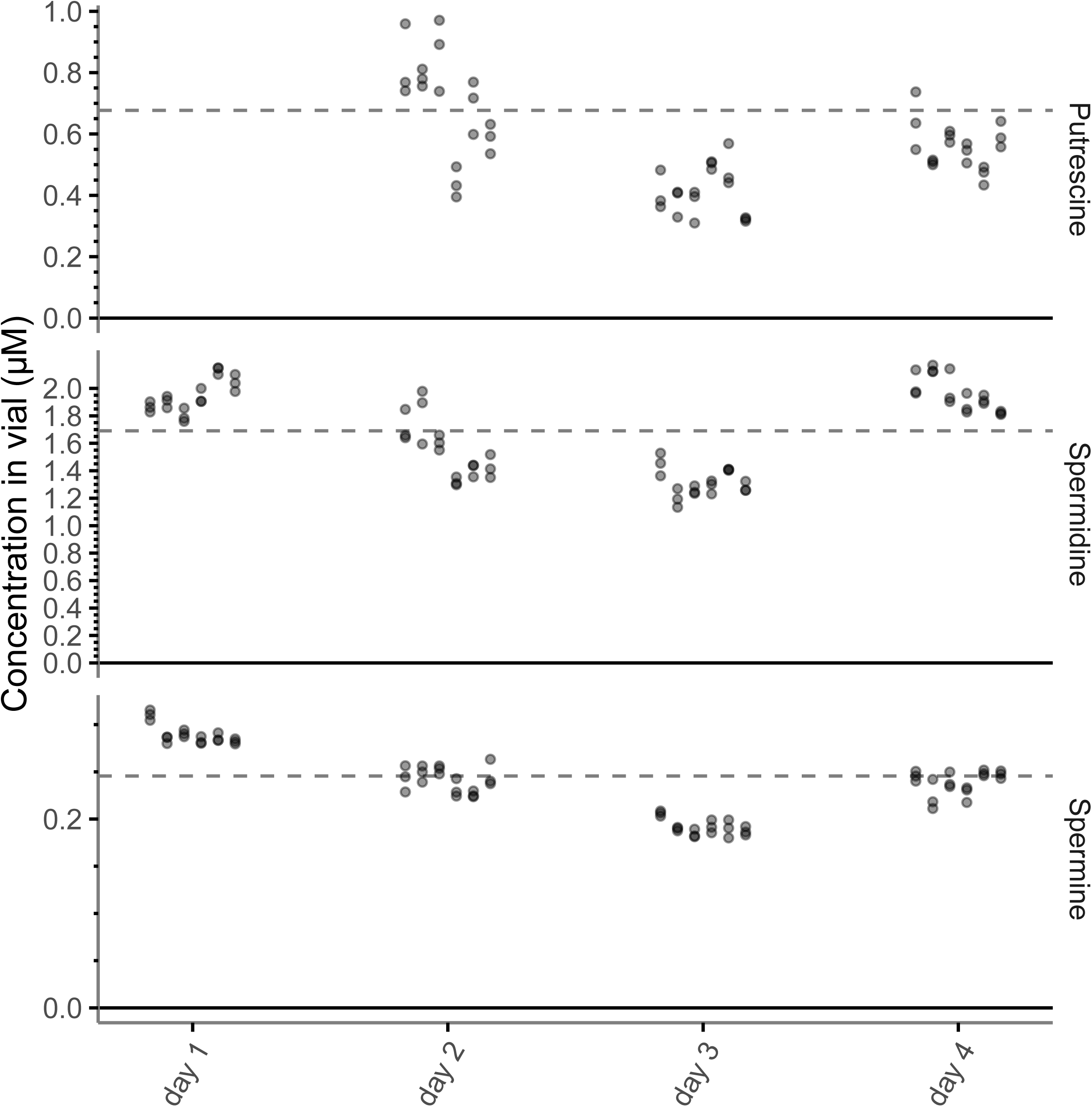
Measured concentrations in cell cultures over four separate days. Technical, inter- and intra-day variation can be inferred from these data. Each “triplet” of points per day and analyte represents triplicate sample measurements. Six cell samples, extracted from six-well culture plates, per day were measured for four non-consecutive days. The dotted lines represent the grand mean overall measurements per analyte.

To determine assay stability, six samples (from a six-well culture plate), equivalent to 500k cells per well. Fresh samples were measured over four quasi-consecutive days. Each sample was injected three times. Sample concentrations for the analytes were determined with calibration curves. Data for Put were missing for day 1 since they were almost completely suppressed by the addition of cadaverine, as mentioned earlier. By validating the assay in this way, we could capture all variability which allowed us to calculate minimal detectable effects (MDEs) that would be significant in standard null-hypothesis significant tests.

Three types of assay variability could be determined, namely inter-day, intra-day and measurement variability. We could distinguish the various types of variation by using a Bayesian multilevel model. This model accounts for hierarchical effects of runs (days) and samples using random effects. This means that it hierarchically models variability across runs and samples, thereby distinguishing the different types of variation. The variation measured between the runs represents the inter-day variability, the variation between samples can be interpreted as the intra-day variability. The remaining variability, *i.e.* variation between the triplicate injections represents measurement variability. Variability is represented as coefficients of variance in percentages (%CV).

As for the sources of variation captured by the different measures, inter-day variation considers sources like variation between culture plates (storage time, incubator position) and differences between calibration curves. Inter-day variation could be, for a big part, interpreted as biological variation since signals were corrected by MFC-normalization for signal drift and variation in extraction and biological material. Measurement variability is mostly due to variation in detector response.

Inter-day variation was lowest for Put with a median value of around 11%, while Spd and Spm were somewhat higher with median values of 18% and 16% respectively (Fig. 7A, Table 5). As can be seen in Fig. 7A, inter-day estimates showed high variability. This is because only four days were compared, and therefore the uncertainty in the estimates is high. Intra-day variation was highest for Put with a median of 16% while those of Spd and Spm were much lower with median values of 7% and 3.6%. Furthermore, the inter-day estimates were quite accurate for Spd and Spm while the uncertainty in the estimate of Put was higher. The remaining measurement error (variation between triplicates) was also highest for Put with about 9.5%, while those of Spd and Spm were under 4%.

**Figure 7.**
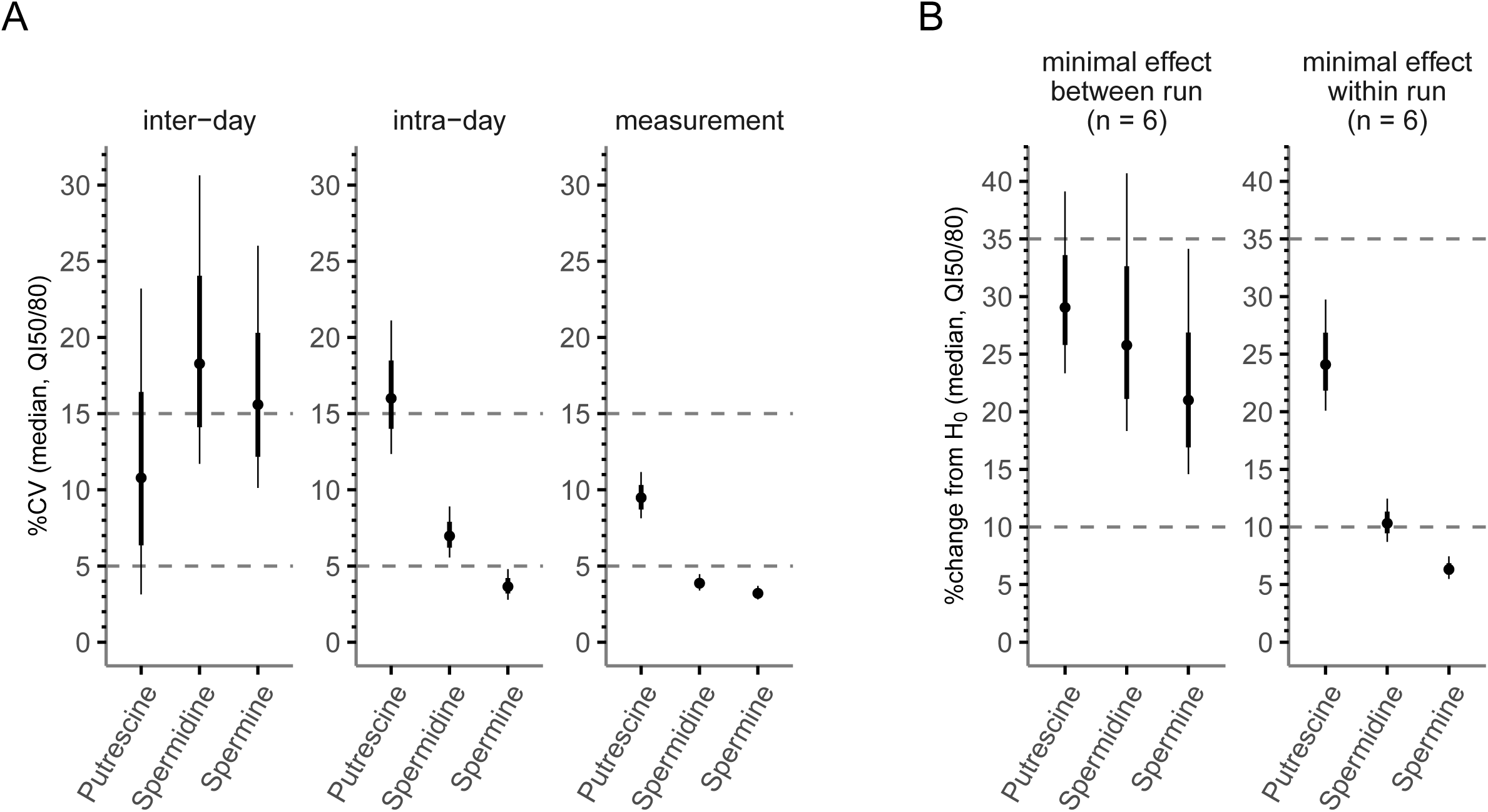
Inter-day, intra-day and measurement variability (A) and minimal significant effects (B) for the assay based on the measured concentrations over four days. Minimal effects were based on pooled standard deviations of inter-, intra-day and measurement (between run) or intra-day and measurement (within run). These pooled standard deviations were then used to calculate a minimal effect that would be significant in a conventional null-hypothesis (H_0_) significance test given α = 0.05 and n = 6 per group.

**Table 5.** Assay metrics including variation and minimal effects. Values are given as the posterior median with the 50% quantile interval between parentheses.

| Analyte | Sources of variation (%CV)<sup>a</sup> |  |  | Minimal effects<sup>b</sup> (% H<sub>0</sub>) |  |
| --- | --- | --- | --- | --- | --- |
|  | inter-day | intra-day | measurement | Between days | Within days |
| Putrescine | 11 | 16 | 9.5 | 29 | 24 |
|  | (6.4-16) | (14-18) | (8.7-10) | (26-34) | (22-27) |
| Spermidine | 18 | 7.0 | 3.9 | 26 | 10 |
|  | (14-24) | (6.2-7.9) | (3.6-4.2) | (21-33) | (9.5-11) |
| Spermine | 16 | 3.6 | 3.2 | 21 | 6.3 |
|  | (12-20) | (3.2-4.2) | (3-3.5) | (17-27) | (5.9-6.8) |

a) %CV: coefficient of variation: 100\*sd/mean.
b) Based on pooled standard deviations. Assuming a two-tailed T-test with $\alpha = 0.05$ and $n = 6$ per group.

By pooling the variances, it was possible to calculate MDEs that would be significant in a null-hypothesis significance test, assuming a two-tailed T-test with α = 0.05 (Fig. 7B, Table 5). If assays need to be compared between days the median of the effects, relative to H_0_, that would be picked up as significant were 44% for Put, 39% for Spd and 32% for Spm. For example, if the mean value in a control group for Spm was 100, the effect being picked up as significant (P < 0.05) would be smaller the 68 or bigger than 132. Sensitivity within runs was much higher, with MDEs for Put of 37%, 16% for Spd and 9.6% for Spm, relative to H_0_. Taking in account these differences in MDEs between and within runs, it is advisable to design experiments so that relevant comparisons are made between groups with the same run.

### Method applications

After evaluating the method, it was used in two separate use cases. First the effect of SAM486A, a well-known inhibitor of the enzyme S-adenosylmethionine decarboxylase (SAMDC) [12], on PA synthesis in DU145 cells was determined. The second use case was to measure intracellular PA levels of DU145 cells incubated in different culture media conditions.

As expected, SAM486A inhibited the synthesis of Spd and Spm completely as can be inferred from the nearly total absence of ^13^C_3_-Spd, ^13^C_3_-Spm and ^13^C_6_-Spm ([M+3], [M+6]) in cells incubated with the inhibitor (Fig. 8). There was also a 2-fold drop in unlabelled Spd and Spm species ([M+0]) which in part can be explained by the fact that the culture medium also contained non-labelled methionine. Moreover, PA pools are not static since PA species are converted to their acetylated forms, can be oxidized and can be excreted from the cell. Therefore, inhibition in synthesis makes that the pools do not get replenished and thus diminish via downstream metabolic processes. On the other hand, Put levels were increased by inhibition of SAMDC. This also makes sense since the SAMDC product, decarboxylated S-adenosylmethionine (dcSAM) is used to convert Put to Spd. Since the production of dcSAMe is strongly inhibited, Put is not converted to upstream PAs and thus accumulates, raising its concentration by about 4-fold.

**Figure 8.**
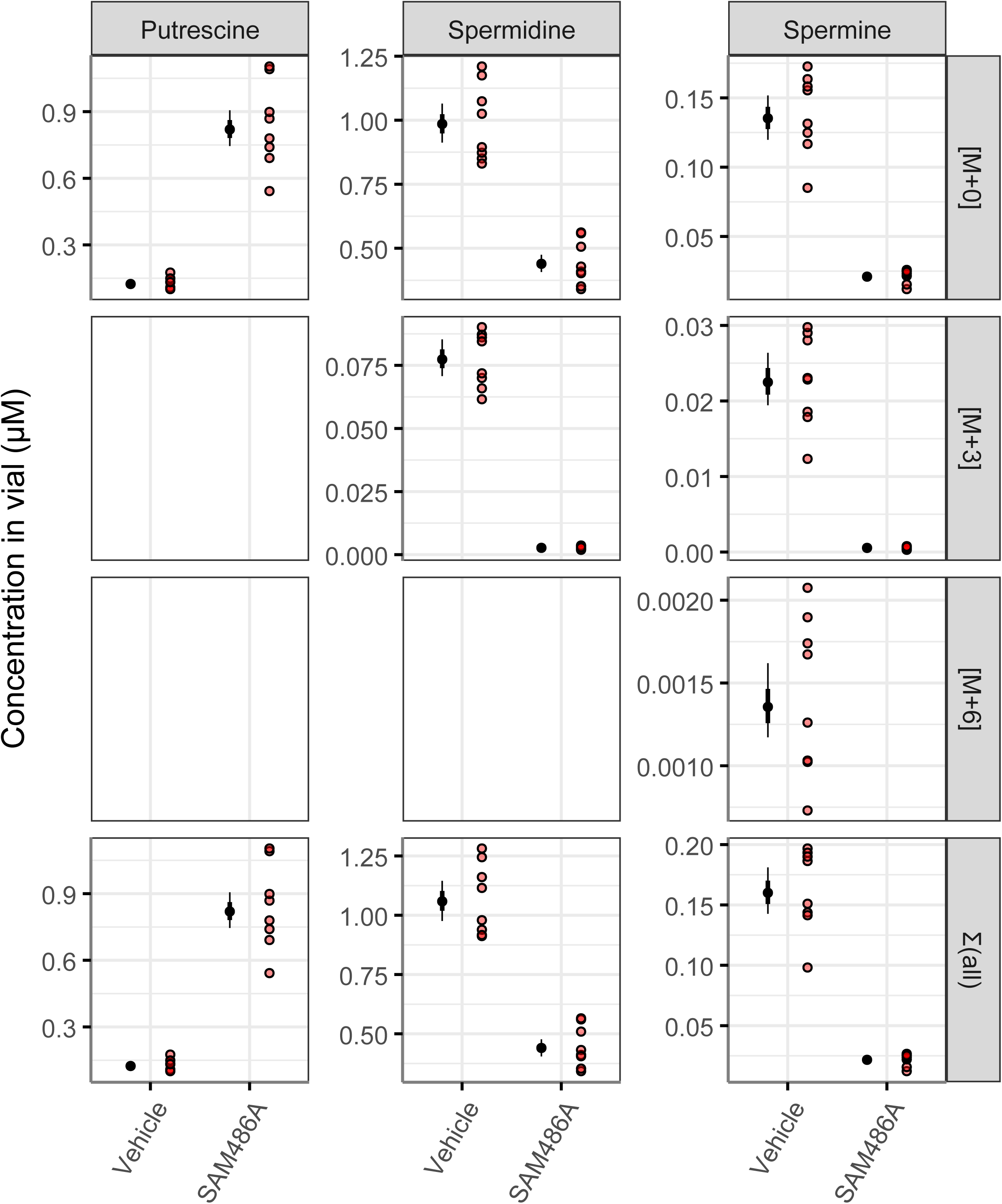
Inhibition of PA synthesis by SAM486A using 13C5 labelled methionine as a precursor. Indicated are the stable labelled PA species for Spd and Spm. [M+3] and [M+6] represent species with ether 3 or 6 ^13^C atoms incorporated [M+0] represent the unlabelled species. Σ(all) is the sum of the concentrations of the labelled and unlabelled species per sample per analyte. Indicated are the group estimates (median in 50/80 quantile intervals, black dots and lines) and individual sample concentrations (red dots).

Next the influence of different culture media types on intracellular PA concentrations was explored. We found that PA concentrations indeed varied widely between culture media and show opposite effects between media and analytes (Fig. 9). When compared to complete DMEM, Put and Spm were elevated in DMEM supplemented with dialyzed serumwhile they were on the same level in HPLM. However, Spm was decreased in dialyzed DMEM while higher in HPLM medium. Assuming no PA presence in media (as it is not an ingredient) Put and Spm synthesis are increased in dialyzed DMEM while Spm synthesis is downregulated.

**Figure 9.**
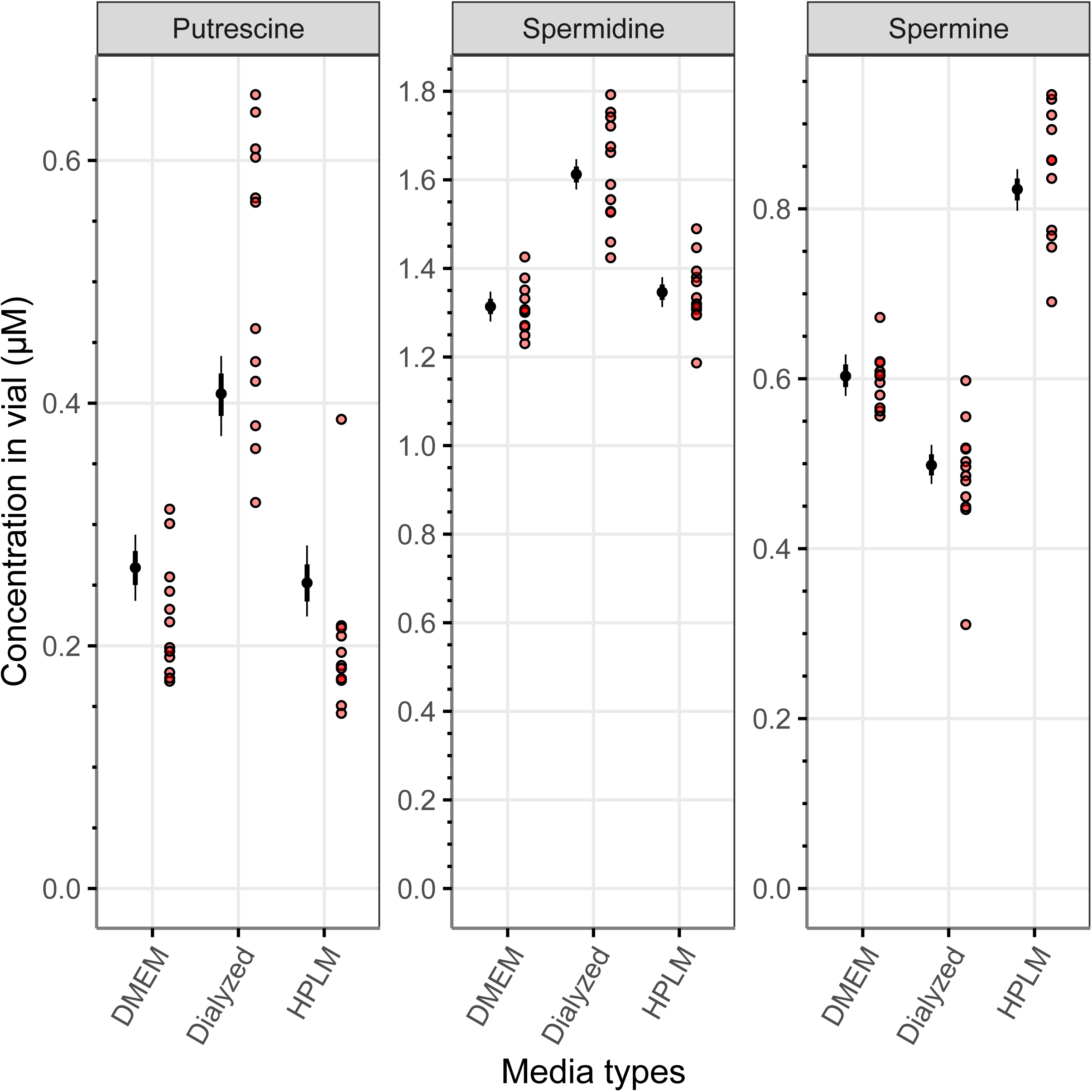
Comparison of cell concentrations of putrescine, spermidine and spermine when using different culture media. Indicated are the group estimates (median in 50/80 quantile intervals, black dots and lines) and individual sample concentrations (red dots).

On the other hand, Spm synthesis is upregulated in HPLM. The biological mechanisms driving these differences are beyond the scope of this study and were not further investigated. However, this finding highlights the relevance of controlling the conditions to analyze the levels of PAs in response to a determined stimulus or the status of the cells in a determined pathology.

## Conclusion

The method described for the analysis of Put, Spm and Spd showed good performance with respect to chromatography and quantification. Moreover, the sample preparation is straightforward and run time is short, making the method ideal for routine use with substantial sample throughput. Furthermore, with the spiked QC comparisons it was possible to estimate signal attenuation due to matrix effects. This allowed us to use a calibration curve in solution, thereby lowering the LLOQ and subsequently correct for the signal suppression of the various analytes. The MDEs calculated based on assay variability are a guideline for the user to adapt the method for testing their own hypotheses. Moreover, preliminary tests (not shown) suggested that this method was also suitable for measuring acetylated polyamines, however some additional tests need to be performed to ensure this. Similarly, the method could likely be applied to other biological matrices, but this also needs additional optimalization steps.

## Supplementary information

Code and pretreated signal data for chromatograms and validation experiments is available in the following repository: https://github.com/smvanliempd/polyamine_analysis.

